# Subtype-Resolved Pain-Signaling Architectures Reveal Conserved Drug-Target Interaction Networks in DRG Nociceptors

**DOI:** 10.64898/2026.04.14.718550

**Authors:** Alexandre Martins do Nascimento, Felipe Monteleone Vieceli, Chao Yun Irene Yan, Eduardo Moraes Reis, Deborah Schechtman

**Affiliations:** Department of Biochemistry, Institute of Chemistry, University of São Paulo, São Paulo, Brazil; Department of Cell and Developmental Biology, Institute of Biomedical Sciences, University of São Paulo, Brazil

**Author notes:** correspondence should be addressed to: D.S. - Institute of Chemistry, 748, Lineu Prestes Av., Butantã, São Paulo, Brazil, 05508-000. (lead contact); Co-corresponding authors: F.M.V. -; A.M.d.N. these authors contributed equally (Alexandre Martins do Nascimento, Felipe Monteleone Vieceli).

**Keywords:** Pain, Analgesics, Drug Targets, Protein-protein Interactions, Signal Transduction, Dorsal Root Ganglia (DRG)

## Abstract

Pain management has been challenging and a major obstacle lies in the limited translational success between preclinical studies, often based on rodent models and evoked nociception behavioral assays, whose validity is often questioned. The dorsal root ganglia (DRG) contains diverse nociceptor subtypes that serve as the primary afferent pathways for detecting painful stimuli and analgesics often target proteins expressed in nociceptors. This makes the distinct protein repertoires and molecular interactors within nociceptor subtypes a key focus for understanding which molecular players drive pain processing and how they may be therapeutically targeted. The confirmation of cross-species conservation of pain-related signaling pathways, mediated by nociceptors, could help to elucidate the molecular mechanisms by which the drugs act across species. In this context, we constructed and compared experimentally-validated protein-protein interaction (PPI) networks based on drug targets and their direct binding partners for nociceptor subtypes supported by single-nuclei transcriptome data from mouse and human DRGs. We found that overall gene expression is more conserved across mice than in human nociceptor subtypes, indicating a higher degree of molecular specialization of human nociceptors. Overall signaling network analyses revealed subtype- and species-specific conservation related to pain signaling, with some particularities, in which key drug targets mediate broader cellular processes beyond pain signaling and neuronal depolarization. Altogether, this resource may help to further understand the molecular mechanisms of specific drug targeting, and the proposed workflow can be used to identify and prioritize pain-related pathways in the DRG, advancing target identification and translational medicine.

## INTRODUCTION

Pain is a multifaceted condition that can be broadly classified as nociceptive, neuropathic, nociplastic or mixed, based on its underlying biological mechanisms, and as acute or chronic according to its duration *(1, 2)*. Currently available analgesics, such as nonsteroidal anti-inflammatory drugs and opioids, are effective in alleviating several forms of nociceptive and acute pain, largely due to the typically self-resolving nature of these conditions *(3, 4)*. In contrast, neuropathic and chronic pain remain considerably more difficult to manage, mainly due to the complex interplay of genetic and environmental factors underlying these conditions, the development of tolerance that often reduces therapeutic efficacy *(5)* and unwanted side effects of some drugs.

It has been proposed that pharmacological targeting of nociceptor-specific proteins and signaling pathways, could lead to improved selectivity *(6)*. Dorsal Root Ganglia (DRG) nociceptors are the primary afferent pathways that encode pain sensation, and key proteins that mediate pain signaling within nociceptor subtypes have been widely explored as pharmacological targets for treating different types of pain including voltage-gated ion channels and cytokines/growth factor receptors *(7)*.

Given the growing availability of high-throughput *omics* datasets related to pain biology, integrative analyses and network-based approaches have emerged as valuable strategies for guiding target identification and drug design *(8)*. We recently proposed that targeting key protein–protein interactions (PPIs) within DRG nociceptor signaling networks could enable the development of analgesics with improved specificity and fewer side effects, and identified promising candidate targets for pain mitigation *(7)*. Here, we use a harmonized atlas, built on single cell and single-nuclei RNA-seq of human and mammalian datasets *(9)*, to perform a direct comparison of cell types across species, as a base to build subtype-specific nociceptive PPI networks. These networks can contribute to further understanding molecular and cellular events mediated by pain targets, which have not yet been addressed.

We applied an analytic workflow to assess whether target expression and associated signaling pathways are conserved across mouse and human homologous nociceptor subtypes. On the side of conservation, we found that signaling network architectures centered on pain targets are largely preserved across both species and nociceptor subtypes. Subtype transcriptional programs show greater conservation among subtypes in mice than in humans. Further, subtype- and species-specific network particularities were identified that may shape the physiology of distinct neuronal subtypes and ultimately influence drug efficacy.

## RESULTS

### Nociceptor transcriptional profile is more conserved in mice than in humans

To compare gene expression profiles of nociceptor subtypes from intra and across different species, we used data from a previously published harmonized RNA-seq atlas of DRG neurons *(9)*. To minimize technical and batch effects, we used a data subset from the harmonized atlas including only nuclei sequenced using the 10x Genomics 3’ RNA platform *(10, 11)*. The data subset used in subsequent analysis include only C-fiber neuronal populations from mouse, guinea pig, cynomolgus macaque, and human datasets (S1A-C; Table S1).

To test if the differences in proportions of nuclei in the different subtypes were significant and could affect data interpretation, we built a contingency table with the proportions of total nuclei counts (excluding mouse-specific subtypes), and performed chi-square tests. The null hypothesis of similar distributions was discarded as we observed significant differences between all four species (Table S2) and between human and mouse only (Table S3). Considering that these variabilities could reflect biological differences in nociceptor subtypes distribution among species, we compared the similarity of gene expression profiles across nociceptor subtypes by evaluating all genes detected in at least 30% of cells in each neuronal subtype (Data file S1). We found that mouse nociceptor subtypes share a highly cohesive global transcriptional profile (mean: 0.718) (Fig. 1A). Human nociceptors exhibited more pronounced global transcriptional variation, indicated by lower inter-subtype similarity (mean: 0.511) (Fig. 1A), while guinea pig and macaque displayed intermediate levels (means 0.651 and 0.641, respectively) (S1D; Data file S2, sheet 1). Of note, the number of nuclei in macaque is greater than mouse, however, this does not reflect a greater similarity across nociceptor subtypes in this species (Fig. S1A), suggesting that the differences are not due to differences in data sampling.

**Figure 1:**
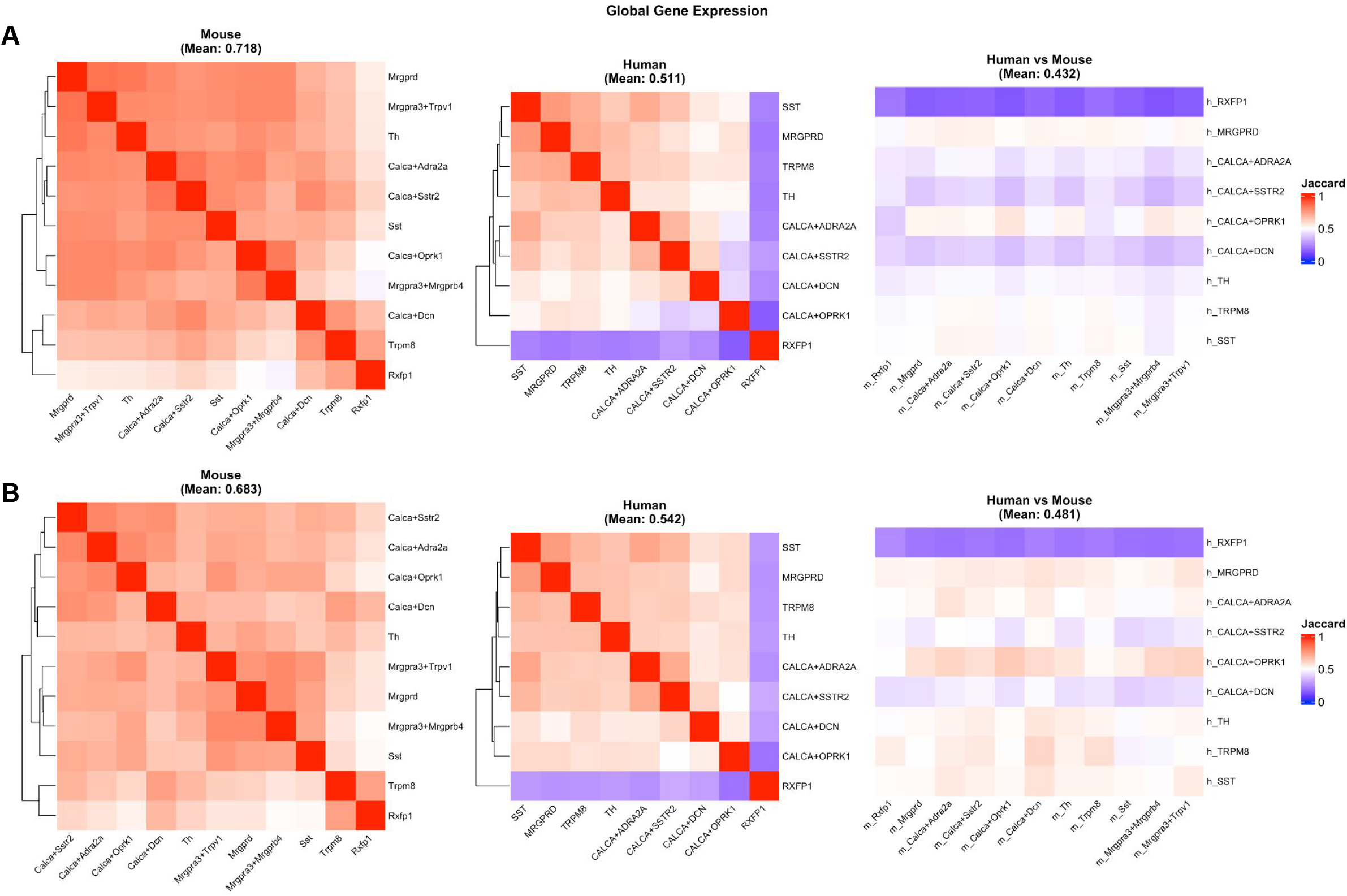
Intra/cross-species gene expression comparison in C-fiber neuronal subtypes. **(A)** Jaccard index heatmaps indicating the similarity of global gene expression (genes detected in at least 30% of cells per cluster) for intra-species comparisons within mouse (mean: 0.718) and human (mean: 0.511), as well as the inter-species comparison between human and mouse (mean: 0.432). **(B)** Jaccard index heatmaps indicating the similarity of pain-related gene expression for intra-species and inter-species C-fiber neuron subtype comparisons, highlighting functional identity functional identity variance within mouse (mean: 0.683), human (mean: 0.542), and between human and mouse (mean: 0.481).

This species-specific transcriptional compartmentalization patterns persisted when only genes associated with pain signaling were considered. Utilizing a curated list of established pain drug targets (Data file S3, sheet 1), we found that the transcription profile similarity across mouse nociceptor subtypes remained highly cohesive (mean: 0.683), yet are distinctly compartmentalized in humans (mean: 0.542) (Fig. 1C). Similarly, guinea pig and macaque pain-related gene expression showed intermediate cohesion (means: 0.603 and 0.614, respectively) (Fig. S1E).

Direct cross-species comparisons indicated that homologous nociceptor subtypes, as classified by Bhuiyan *et al. (9)*, between humans and mice share only moderate transcriptional similarity, both globally (mean: 0.432) and regarding pain-related (mean: 0.481) gene expression (Fig. S1D-E; Data file S2, sheet 2). The observation of comparable intra/cross-species Jaccard index values using all genes (Fig. 1A) or only pain-related genes (Fig. 1B) suggests that these patterns reflect similarities/differences in the structure and function of pain networks across neuron subtypes and species. We posited that the transcriptional divergences observed between mouse and human nociceptor neurons may reflect functional differences in pain-related drug targets, which could explain failures of preclinical rodent studies to yield effective clinical therapies. Thus, mapping subtype-specific protein-protein interaction networks might further support prioritization of new analgesic targets.

### Pain signaling networks are conserved between homolog nociceptor subtypes

To address the possible pain signaling rewiring in nociceptor subtypes, we analyzed PPI networks involving pain targets and their binding partners *(7)*. For this, the list of expressed genes from the previous section was used to build networks of experimentally validated PPIs for each nociceptor subtype using the STRING database *(12, 13)*. For each network, curated pain drug targets (Data file S3, sheet 1) were mapped and their first interaction layer was preserved for further analysis (Fig. 2A). Thus, networks considering pain drug targets as hubs and the interaction of the direct binding partners as edges were built for each nociceptor subtype.

**Figure 2:**
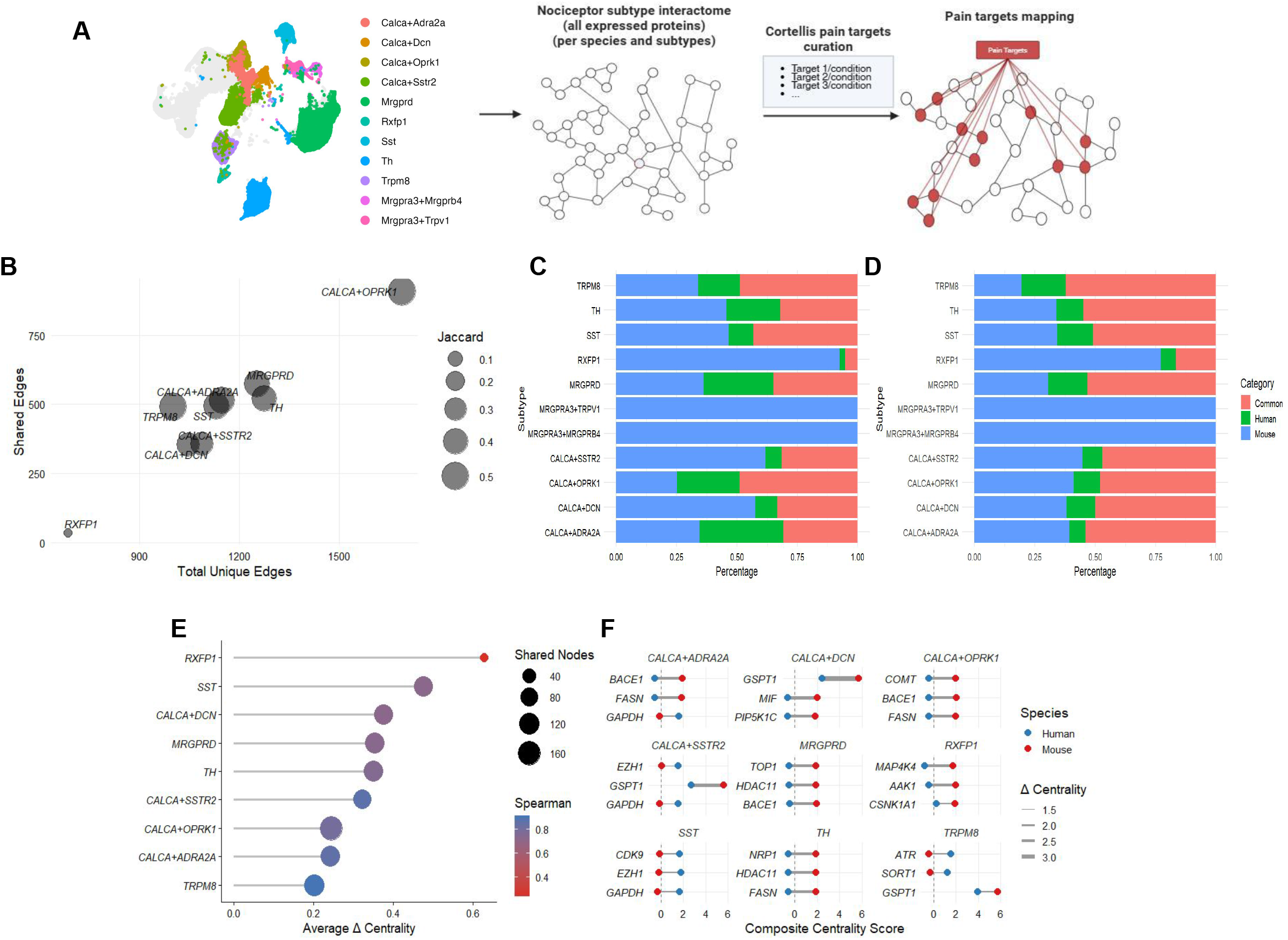
Pain signaling networks conservation analysis between human and mouse nociceptors. **(A)** workflow of how signaling networks were obtained: lists of genes expressed in at least 30% of nociceptor clusters nuclei were used to build protein-protein interaction networks. A curated list of pain molecular targets was used to map the pain hubs within the networks, to identify direct binding partners. **(B)** similarity index of first-layer interactions from shared pain target hubs, indicating high conservation amongst subtypes. **(C)** Percentage of edge distribution amongst nociceptor clusters. **(D)** Percentage of hub distribution amongst nociceptor clusters. **(E)** Conservation of shared pain target hubs amongst homolog nociceptors subtypes, indicating high conservation amongst subtypes. **(F)** Centrality shifts of the three more divergent hubs from each subtype assessed corroborate high overall conservancy.

Topological analysis of pain-related genes shared between homolog subtypes of nociceptors revealed that there is first-layer edge conservation amongst clusters, reflecting preservation of drug targets signaling context within these nociceptor subtypes (Fig. 2B). Notably, two subtypes do not cluster with the others: *CALCA+OPRK1*, which has the highest number of edges (shared and unique), but has a high conservation index; *RXFP1*, treated here as a special case due to low counts of hubs available for humans (Fig. 2B). This network conservation analysis revealed that common hubs across homologous nociceptor subtypes share an average of 50% similarity, while common edges share only 25% similarity. Additionally, the network comparison revealed more mouse-specific expressed hubs (pain drug targets) and edges (direct binding partners) than human-specific ones (Fig. 2C-D, Data File S3, sheet 2).

With regards to hubs, we found conserved centrality indexes amongst mouse and human homolog nociceptors (approximately > 70% correlation on average) (Fig. 2E). Inspection of the top three most divergent hubs in each nociceptor subtype revealed no major centrality shifts between species, further confirming the overall network topology conservation (Fig. 2F, Data File S3, sheet 3). Thus, shared pain drug targets hubs tend to have conserved network structures between species, directly interacting with other protein edges with few shifts.

### Signaling network analysis of pain hubs reveals conservation between shared genes and rewiring for species-specific genes

We then performed differential network analysis to address subnetwork rewiring across species (in their nodes and first-layer interactors) (Data File S3, sheet 4-5). We found that an average of 25% of the subnetworks are identical between species in any homologous nociceptor subtype, while an average of 45% of subnetworks are modified, and 30% represent species-specific networks (Fig. 3A). Since a gain/loss of a single component could classify the subnetwork as ‘modified’, we quantified the similarity between subnetworks by analyzing the overlap between the networks with the Jaccard similarity index. This analysis revealed a comparable distribution indexes between human and mouse pain gene networks, predominantly with over 50% identity (Fig. 3B), suggesting conservation of signaling architecture across all nociceptor subtypes, with exception of *RXFP1*. The top three most divergent subnetworks of each neuronal subtype revealed that, overall, there is a tendency of human nociceptors for losing direct binding partners. This occurs frequently among the same hubs, such as *GSPT1, CNSK1A1* and *VCP* (Fig. 3C).

**Figure 3:**
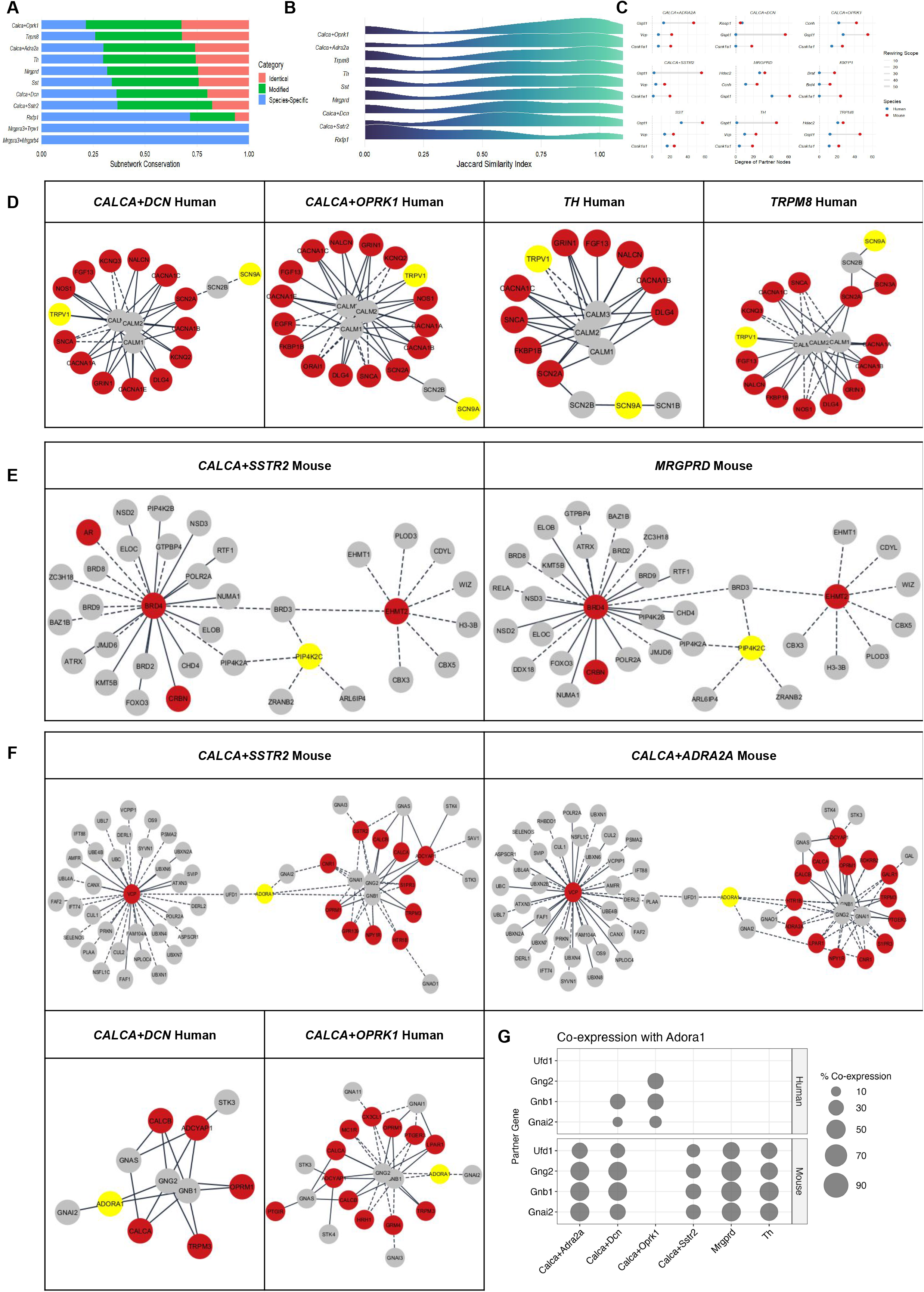
Pain subnetworks signaling rewiring analysis. **(A)** Percentage of identical, modified and species specific subnetworks in nociceptor subtypes. **(B)** Density distribution of Jaccard similarity index for subnetworks. **(C)** Rewiring scope (delta number of edges) of human nociceptors, taking into consideration the three most divergent subnetworks of each subtype. Density of 1 is related to identical subnetworks, while density of 0 is related to species-specific subnetworks. **(D)** *SCN9A*, **(E)** *PIP4K2C*, **(F)** *ADORA1* expanded signaling networks (+ 4 interaction layers). Pain hubs = yellow; Pain targets from curated list = red; First layer interactors mapped = gray. Solid line = common interaction (mice/humans); Dashed line = species-specific interaction. **(G)** Co-expression of *ADORA1* with its direct interactors in *CALCA+SSTR2, CALCA+ADRA2A, CALCA+DCN* and *CALCA+OPRK1* nociceptor subtypes.

Sodium voltage-gated ion channel 1.7 (Na_V_1.7), encoded by *SCN9A*, a major clinical target for pain management *(14)*, is shared in 8 out of 9 homolog subtypes, with negligible centrality shift among different networks (Fig. S2A), even when it interacted with more than one binding partner, as in *TH* nociceptors. We expanded its subnetwork, and found a conserved signaling architecture between different subtypes of both mice and human nociceptors, as it is connected to multiple ion channels signaling cascades and other pain targets via *SCN2A/B* subunits (Fig. 3D). A similar context for *TRPV1*, another very explored channel in multiple pain types *(15)*, was found in the network (shared in 6 out of 9 homolog subtypes; Fig. S2B).

The connection between *TRPV1* and *SCN9A* subnetworks is absent in mouse *TH* and *CALCA+DCN* nociceptors (Fig. 3D), that do not express one of the channels, thus targeting either of these two proteins could perturb both networks but not in these neurons in mice.

We also investigated hubs categorized as mouse-specific based on single nucleus RNAseq expression to find signaling pathways that are absent in human homolog subtypes. The drug target, *PIP4K2C* (Phosphatidylinositol-5-phosphate 4-kinase gamma) is a regulator of PIP2 levels, critical for nociceptive signaling *(16, 17)* that is not expressed in human nociceptors. In mouse *CALCA+SSTR2* and *MRGPRD* subtypes it also interacts with chromatin remodeling proteins such as *BRD3/4 (18)*, that in turn interacts with *EHMT2*, another epigenetic modulator that is a drug target, representing mouse-specific signaling pathways in these specific nociceptors that are absent in humans (Fig. 3E). A drug that targets both *PIP4K2C* and *PIP5K1C* lipid kinases has been launched (Data file S3, sheet 1) *(16)*. While *PIP4K2C* is only expressed in mouse nociceptors, *PIP5K1C* is found in both mice and human nociceptors (Data file S3, sheet 2).

Adenosine A1 Receptor (ADORA1) is a GPCR-coupled receptor that regulates neuron calcium influx, contributing to pain signaling *(19)*. This receptor was a common target for migraine via combination of ergotamine tartrate/caffeine *(20)*. However, the drug was discontinued because of its many adverse effects arising from its activity in the central nervous system *(21)*. This target is predominantly mouse-specific based on expression (Data file S3, Sheet 2), but it is expressed in both species in *CALCA+DCN* neurons, and is present only in the human *CALCA+OPRK1* nociceptors. We took advantage of this particular expression profile to address potential signaling pathways that differ between species based on presence/abscence of the hub’s direct interactors.

We found conserved GPCR signaling in all subnetworks, but a rewiring in the link between *ADORA1* signaling and *UFD1-VCP* proteolytic system (Fig. 3F) in mouse *CALCA+SSTR2, CALCA+ADRA2A* and *MRGPRD* nociceptors. We further confirmed this finding at the single-nucleus level by assessing the specific co-expression of *ADORA1* with its first layer interactors (Fig. 3G) including *UFD1* that was not expressed in the same nociceptor subtype or any other subtype in humans (Data file S3, sheet 6). Taken together, this analysis suggests that this new interaction can in fact occur in these neuronal subtypes and that this drug acts on a mouse-specific pathway that is not necessarily related to pain. This finding suggests that, even with elevated overall signaling conservancy, some cases may represent major signaling rewiring between species in homologous nociceptors.

Even though the human data has low counts for specific nociceptor subtypes (e.g. *CALCA+OPRK1*, Table S1), affecting distribution (Table S2-3), we were able to find human-specific pairs expression in underrepresented subtypes.

## DISCUSSION

Targeting PPIs is a way of inhibiting a specific signaling pathway and this is an emerging and promising strategy, with growing evidence of clinical success for other conditions *(22)*. Previously, we suggested that selection of PPIs that can be targetable in nociceptors could be obtained by integrating nociceptor expression data with network analysis *(7)*. Here we used single-nuclei RNAseq data from DRG nociceptor neurons *(9–11)* to infer subtype-specific PPIs comprising tested pain drug targets and their experimentally validated binding partners in mice and humans and perform topological network analyses to verify their conservation between species.

Considering that the harmonized DRG atlas clustering was built based on marker gene expression, but not entirely considering physiology due to the fact that the function of some nociceptor subtypes are not yet well defined *(9)*, we propose that the integration with PPI networks of pain drug targets, as performed in this study, can support and refine the functional classification related to pain signaling across neuronal subtypes. Here, we found elevated conservation of hubs and edges, reflecting that they likely represent true homologous nociceptor classes with preserved functional roles regarding pain signaling.

Through our workflow, we found conserved signaling topology and shared expression amongst nociceptor subtypes for known and promising pain targets. For instance, AP2 associated kinase (AAK1), which regulates internalization of membrane receptors that induce nociceptor sensitization, has been experiencing a smooth translation from preclinical to clinical models *(23, 24)*. An AAK1 selective inhibitor is currently being developed at phase II clinical trials to treat neuropathic pain, showing promising results regarding antinociception and safety *(25)*. Our dataset reveals that this hub has shared expression amongst all similar subtypes, with centrality and edge conservation. Notably, *NECAP1*, which binds directly to *AAK1* in human *MRGPRD* and *CALCA+OPRK1* nociceptors, further reinforces the clathrin-mediated internalization of ion channels in this network *(26)*.

In contrast, signaling network analysis suggested that some therapeutic targets derived from mouse studies may affect humans differently due to diverging networks in human nociceptor subtypes, as compared to mice. For instance, *PIP4K2C*, which interacts with chromatin remodeling proteins in mice, and *ADORA1* with proteins involved in proteolysis. Cases like *PIP4K2C* and *ADORA1* exemplify species-specific network rewiring, where direct interactors are shifted or absent in human homologs. This rewiring may not directly affect nociceptor-mediated pain processes that are centered in ion channels and neuronal depolarization, but might ultimately affect the outcome of the drug, due to the fact that the targets may act on processes that maintain neuronal homeostasis.

Many predictive tools specific to pain are now available for researchers, as well as high throughput expression data for a large variety of pain conditions in both mice and humans. Additionally, given the urgency of approval of analgesic strategies for chronic pain, it is essential that this information is used to improve the odds of success. Our data suggests that shared expression amongst homologous nociceptors and conserved signaling architecture is an additional factor to consider when addressing a pain target as well as to evaluate what other effects may be expected that can contribute to the overall efficacy of the drug.

A limitation of our study is the fact that we did not consider protein expression and RNA was only considered in terms of nuclei with positive/negative expression within each neuron subtype population. We considered the co-expression between a hub and all its interaction partners (subnetwork) to establish that the inferred signaling networks are active in the same cell. Since large subnetworks in clusters composed of a small number of nuclei could skew the results to < 10% co-expression frequencies, co-expression analysis for each hub-edge pair was also performed.

Additionally, we mapped PPI networks based on expression data from basal states, we have not considered change in pain conditions, which may imply the remodeling of cluster distribution, such as *ATF3* or putative silent nociceptors *(27, 28)*. Expressions in these clusters are pain type-dependent, and dependent on an intricate neuronal network involving both central and peripheral nervous systems that could lead to PPI network remodeling that may also vary with evolution of the pain condition. However, we pinpoint the fact that this remodeling arises from a common background. Another limitation is the currently available physical interaction evidence, which may be underrepresenting some hub interactions in our networks. With advances in AI to predict binding partners *(29, 30)*, and new experimental approaches *(31, 32)*, these networks may grow and help further characterize subtypes of nociceptors in pain signaling and nociceptor subtype physiology.

Furthermore, our analysis is focused on DRG nociceptors, but non-nociceptors and non-neuronal cells are known contributors to pain signaling in the peripheral nervous system *(33, 34)*, and could be explored with a similar workflow, as they can express many of the known pain targets.

Altogether, we propose that the systematic analysis of the structure and conservation of pain-related genes and their interactors in mouse and human nociceptors subtypes may contribute to a deeper comprehension of the signaling pathways mediated by these drug targets, to the understanding of how these pathways contribute to the overall effect of the drugs in different species and possibly help to identify new analgesic targets.

## METHODS

### Dataset Preparation

To characterize the transcriptional profiles of specific sensory neuron populations, previously published single-cell and single-nucleus RNA sequencing (snRNA-seq) data from dorsal root ganglia (DRG) neurons was used *(9)*. Metadata was extracted from the full harmonized dataset to evaluate technical features, including total cell/nuclei counts across different sequencing platforms and species for each neuron subtype. To minimize technical batch effects for the cross-species comparative analyses, the dataset was subsequently filtered to strictly include nuclei sequenced using the 10x 3’ RNA platform.

To address differences in counts distribution, we performed regular chi-squared comparison between all species, and between human and mouse, based on raw counts of sequenced nuclei (Table S1). Mouse-specific subtypes were not included in the analysis.

### Inter/Cross-Species Neuronal Subtype Transcriptional Similarity

Gene expression was established by evaluating normalized count values (greater than zero) for each gene within each C-fiber cluster, using the neuronal subtype annotation retrieved from the original study *(9)*. To ensure robust gene set comparisons, only genes detected in at least 30% of the nuclei within a given subtype were retained for further analyses (Data File S1). To quantify the global transcriptional similarity between nociceptor subtypes, both intra-species and inter-species, Jaccard similarity matrices were computed based on these filtered gene sets. The Jaccard index was calculated as the number of genes expressed in both subtypes (intersection) divided by the total number of genes expressed in either subtype (union). Global intra-species mean conservation was calculated using values from the upper triangle of the similarity matrices (to avoid redundant reciprocal comparisons and exclude self-comparisons), while inter-species divergence (human vs. mouse) was calculated using the entire comparison matrix. To assess functional conservation specifically related to pain signaling, the previously generated gene sets were further filtered using a curated list of pain-related genes (see “Dataset Curation” for details). New intra- and inter-species Jaccard similarity matrices and mean scores were then recalculated exclusively using this filtered subset of pain-associated genes, using the identical intersection-over-union methodology (Data File S2).

### Curation of a Pain-Related Gene Dataset

For molecular pain targets list retrieval, we conducted a search in Clarivate’s Cortellis Drug Discovery Intelligence using: “Pain” (Term); “Genes & Targets” (Knowledge Area); “*Homo sapiens*” (Condition Filter). Search was up to date until March 15, 2026. Targets were filtered to keep unique entries and associated pain conditions, drugs developed for each condition and highest phase of experimental development were registered (Data file S3).

### Network Analysis

The filtered gene sets detected in more than 30% of the nuclei of each cluster were used to create subtype-specific PPI networks using the StringApp *(12, 13)* in Cytoscape (V.3.10.4). The *M. musculus* gene set was ortholog mapped to match *H. sapiens* using g:Profiler g:Orth function *(35)*. To rule out false-negative interactions, PPIs were mapped across species using a matched 0.5 *H. sapiens* experimental evidence score cut-off, with singletons filtering.

Undirected topological analysis was carried out using NetworkAnalyzer (4.5.0) *(36)*. The degree (number of connected edges), betweenness centrality (capacity to mediate information flow) and neighborhood connectivity (average degree of interacting genes) for each node was retrieved. A combined centrality score was calculated for each node by averaging the Z-scores of individual centrality measures, and shared pain nodes across subtype homologs were compared to derive an absolute delta centrality. A Spearman correlation of combined centrality scores was calculated. The first interaction layer, defined as the direct interactors of pain targets (adjacent edges and their connected nodes), was retained. For each subtype, the intersection (shared edges across orthologs) and the total number of edges was calculated to generate a Jaccard similarity index. The same procedure was applied for differential subnetwork analysis (Data File S3).

## Data availability

The authors declare that all the data supporting the findings of this study are contained within the paper and the supplementary materials. Reclustering scripts and Cytoscape network files are available at 10.5281/zenodo.19545165.

## Author contributions

D.S. supervised the research; D.S., F.M.V and A.M.d.N. conceived and designed research; F.M.V. performed expression data analysis. A.M.d.N. performed dataset curation, and network analysis. All authors discussed, wrote, edited and approved the final version of the manuscript.

## Funding

This work was supported by Fundação de Amparo à Pesquisa do Estado de São Paulo (FAPESP): Grant No. 2019/06982-6 (to D.S.), fellowship No. 2024/18174-0 (to A.M.d.N.), fellowship No. 2020/13929-1 (to F.M.V.). Conselho Nacional de Desenvolvimento Científico e Tecnológico (CNPq): fellowship No. 305547/2024-0 (D.S.). This study was also financed in part by the Coordenação de Aperfeiçoamento de Pessoal de Nível Superior - Brasil (CAPES) - Finance Code 001.

## SUPPLEMENTARY MATERIALS

**Table S1:** Nuclei counts of C-fiber clusters Average Silhouette Widths (ASW) within the UMAP space.

| Species | Atlas Annotation | Nuclei Count |
| --- | --- | --- |
| C. Macaque | Calca+Adra2a | 698 |
| C. Macaque | Calca+Dcn | 2 |
| C. Macaque | Calca+Oprk1 | 4 |
| C. Macaque | Calca+Sstr2 | 516 |
| C. Macaque | Mrgprd | 4277 |
| C. Macaque | Rxfp1 | 42 |
| C. Macaque | Sst | 665 |
| C. Macaque | Th | 277 |
| C. Macaque | Trpm8 | 106 |
| Guinea pig | Calca+Adra2a | 186 |
| Guinea pig | Calca+Dcn | 3 |
| Guinea pig | Calca+Oprk1 | 2 |
| Guinea pig | Calca+Sstr2 | 377 |
| Guinea pig | Mrgprd | 933 |
| Guinea pig | Rxfp1 | 27 |
| Guinea pig | Sst | 146 |
| Guinea pig | Th | 368 |
| Guinea pig | Trpm8 | 76 |
| Human | Calca+Adra2a | 60 |
| Human | Calca+Dcn | 10 |
| Human | Calca+Oprk1 | 6 |
| Human | Calca+Sstr2 | 556 |
| Human | Mrgprd | 338 |
| Human | Rxfp1 | 4 |
| Human | Sst | 158 |
| Human | Th | 15 |
| Human | Trpm8 | 57 |
| Mouse | Calca+Adra2a | 294 |
| Mouse | Calca+Dcn | 120 |
| Mouse | Calca+Oprk1 | 350 |
| Mouse | Calca+Sstr2 | 693 |
| Mouse | Mrgpra3+Mrgprb4 | 156 |
| Mouse | Mrgpra3+Trpv1 | 152 |
| Mouse | Mrgprd | 2153 |
| Mouse | Rxfp1 | 115 |
| Mouse | Sst | 438 |
| Mouse | Th | 1512 |
| Mouse | Trpm8 | 477 |

**Table S2:** Chi-square residuals calculated for each subtype considering all species according to the expected distribution, based on nuclei counts (Table S1), extracted from the harmonized dataset (p-value = 0.004998).

| Species | Calca+Adra2a | Calca+Dcn | Calca+Oprk1 | Calca+Sstr2 | Mrgprd | Rxfp1 | Sst | Th | Trpm8 |
| --- | --- | --- | --- | --- | --- | --- | --- | --- | --- |
| Macaque | 8.444 | -7.172 | -11.856 | -12.230 | 19.905 | -3.998 | 3.662 | -20.565 | -10.950 |
| Guinea pig | 1.780 | -3.508 | -6.620 | 5.624 | -2.590 | 0.443 | -2.903 | 4.820 | -1.896 |
| Human | -3.405 | -0.038 | -4.058 | 31.205 | -9.960 | -2.689 | 5.114 | -11.585 | 0.454 |
| Mouse | -8.275 | 9.497 | 17.948 | -4.450 | -14.671 | 5.066 | -4.348 | 23.577 | 12.242 |

**Table S3:** Chi-square residuals calculated for Human and Mouse subtype according to the expected distribution, based on nuclei counts extracted from the harmonized dataset (p-value = 0.004998).

| Species | Calca+Adra2a | Calca+Dcn | Calca+Oprk1 | Calca+Sstr2 | Mrgprd | Rxfp1 | Sst | Th | Trpm8 |
| --- | --- | --- | --- | --- | --- | --- | --- | --- | --- |
| Human | 0.270 | -2.445 | -6.847 | 24.589 | -3.453 | -3.507 | 6.120 | -14.860 | -3.252 |
| Mouse | -0.120 | 1.082 | 3.029 | -10.878 | 1.527 | 1.551 | -2.708 | 6.574 | 1.439 |

**Data File S1:** Expression metrics of genes detected in at least 30% of the cells within each cluster.

**Data File S2: (Sheet 1)** Jaccard similarity matrix indicating the intra-species variation of global gene expression across C-fiber neuronal subtypes. **(Sheet 2)** Jaccard similarity matrix indicating the intra-species variation of pain-related gene expression across C-fiber neuronal subtypes.

**Data File S3: (Sheet 1)** Curated pain targets and clinical overview. **(Sheet 2)** Mapping of pain targets within nociceptor subnetworks. **(Sheet 3)** Normalized network centrality parameters for curated pain targets. **(Sheet 4)** Subnetwork co-expression frequencies. **(Sheet 5)** Subnetwork similarity analysis. **(Sheet 6)** Expression of pain target hubs and their direct interactors in their respective clusters.

**Supplementary Figure 1:**
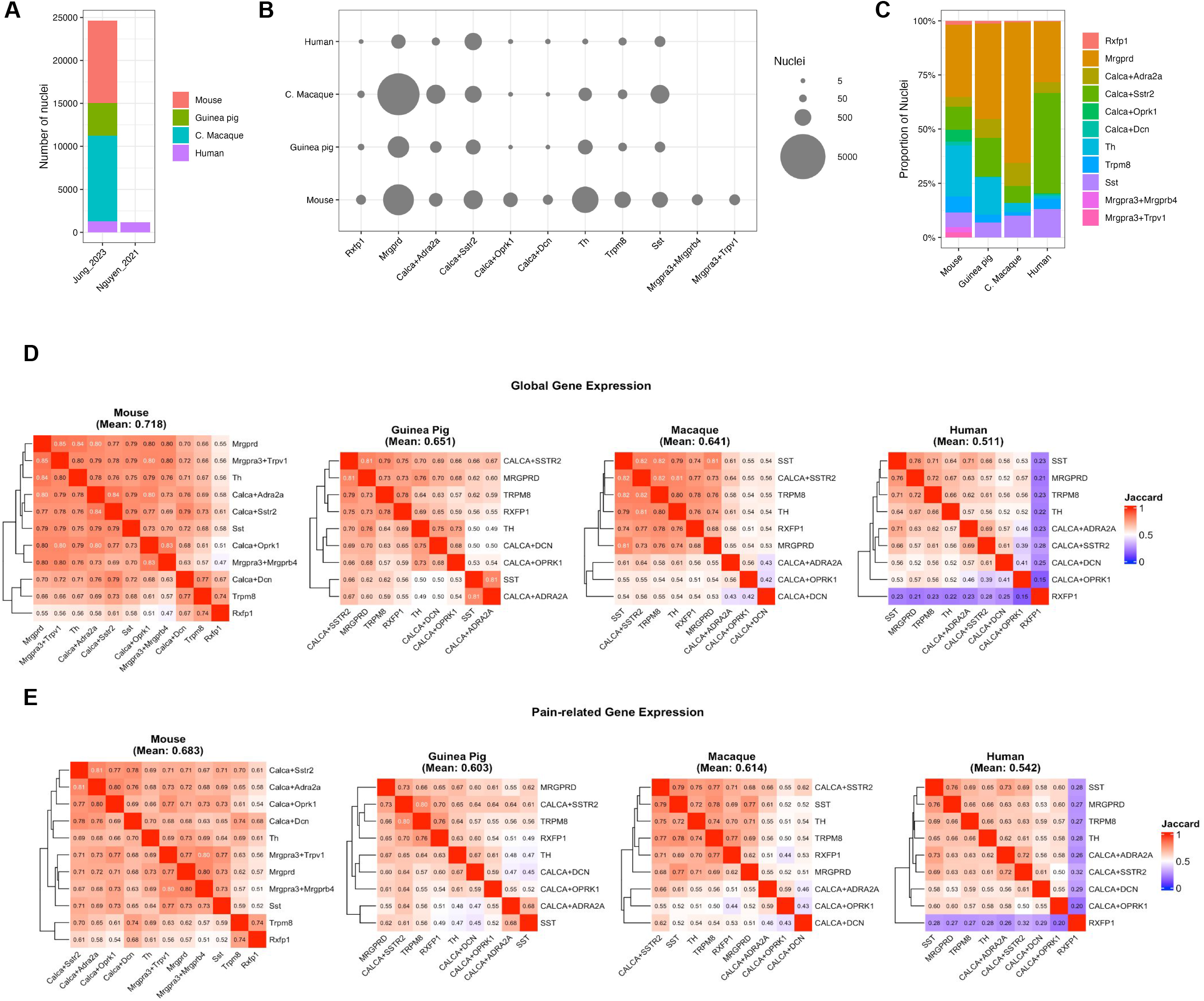
Technical features of the harmonized dataset and intra-species comparison of gene expression in C-fiber neuronal subtypes. **(A)** Bar plot displaying the total nuclei contributed per study using the 10x 3’ RNA platform. **(B)** Nuclei counts amongst species. **(C)** Nuclei proportion per species. **(D)** Jaccard similarity heatmaps indicating the intra-species variation of global gene expression (genes detected in at least 30% of cells per cluster) across all clusters identified in mouse (mean: 0.718), guinea pig (mean: 0.651), cynomolgus macaque (mean: 0.641), and human (mean: 0.511). **(E)** Jaccard similarity heatmaps indicating the intra-species variation of pain-related gene expression: mouse (mean: 0.685), guinea pig (mean: 0.603), cynomolgus macaque (mean: 0.614), and human (mean: 0.543).

**Supplementary Figure 2:**
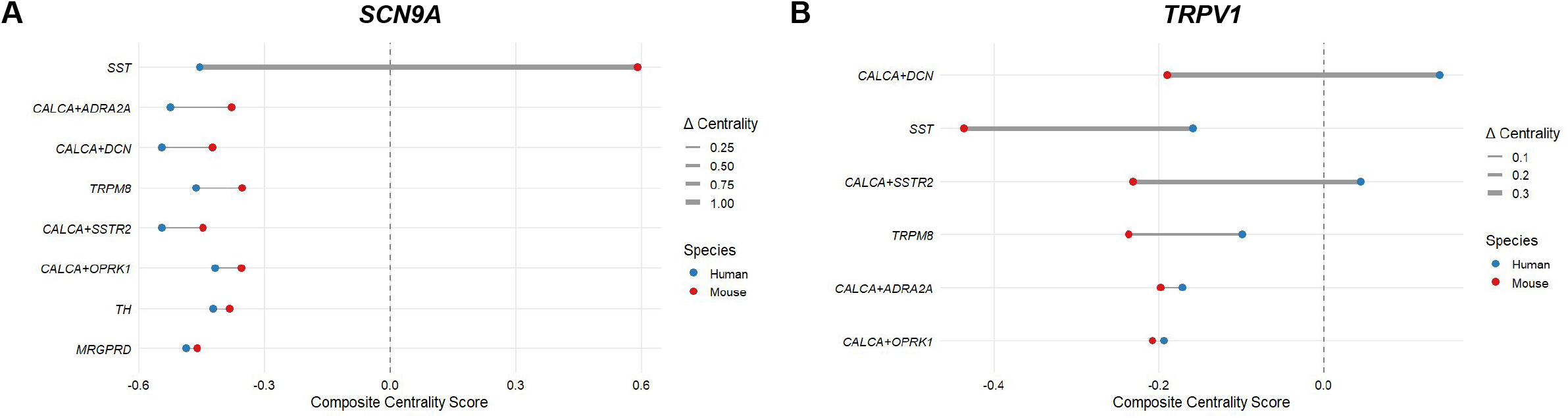
Hubs centrality distribution amongst nociceptor subtypes. **(A)** *SCN9A* and **(B)** *TRPV1* centrality shifts between mouse and humans across subtypes.

## Notes

### Competing Interest Statement

The authors have declared no competing interest.

